# miR-6883 downregulates HIF1α in colorectal and breast cancer cells

**DOI:** 10.1101/2023.09.05.556385

**Authors:** Nicole Jensen-Velez, Lindsey Carlsen, Wafik El-Deiry

## Abstract

Colorectal cancer (CRC) and breast cancer (BC) are deadly diseases that rank as the second and fourth leading causes of cancer-related deaths, respectively. We have previously shown that miR-6883 targets CDK4/6 and that palbociclib-mediated CDK4/6 inhibition destabilizes HIF1α. We hypothesize that miR-6883 downregulates HIF1α in CRC and BC cells. miR-6883 was transfected under normoxia or hypoxia and western blot analysis revealed that miR-6883 downregulates CDK4/6 and HIF1α in CRC and BC cells, pointing to miR-6883 as a promising therapeutic to target hypoxic cancers or HIF1α-deregulated tumor cells. Future studies will further investigate miR-6883 as a cancer biomarker, effects on HIF–related proteins and therapeutic uses *in vivo*.

Figure 1.
miR-6883 downregulates CDK4/6 and HIF1α in colorectal and breast cancer cells
A) Colorectal cancer cells were treated with doses of palbociclib ranging from 0-20 µM. Cell viability was measured by imaging the bioluminescent signal after addition of CellTiter-Glo reagent. B) Percent viability of colorectal cancer cells treated with palbociclib was calculated and nonlinear regression analysis was completed using GraphPad Prism software. C-H) CRC cells were transfected with miR-6883 under normoxia or hypoxia (<0.5% O2) and protein levels of CDK4/6, HIF1α, Glut1, and Ran were measured by Western blot.

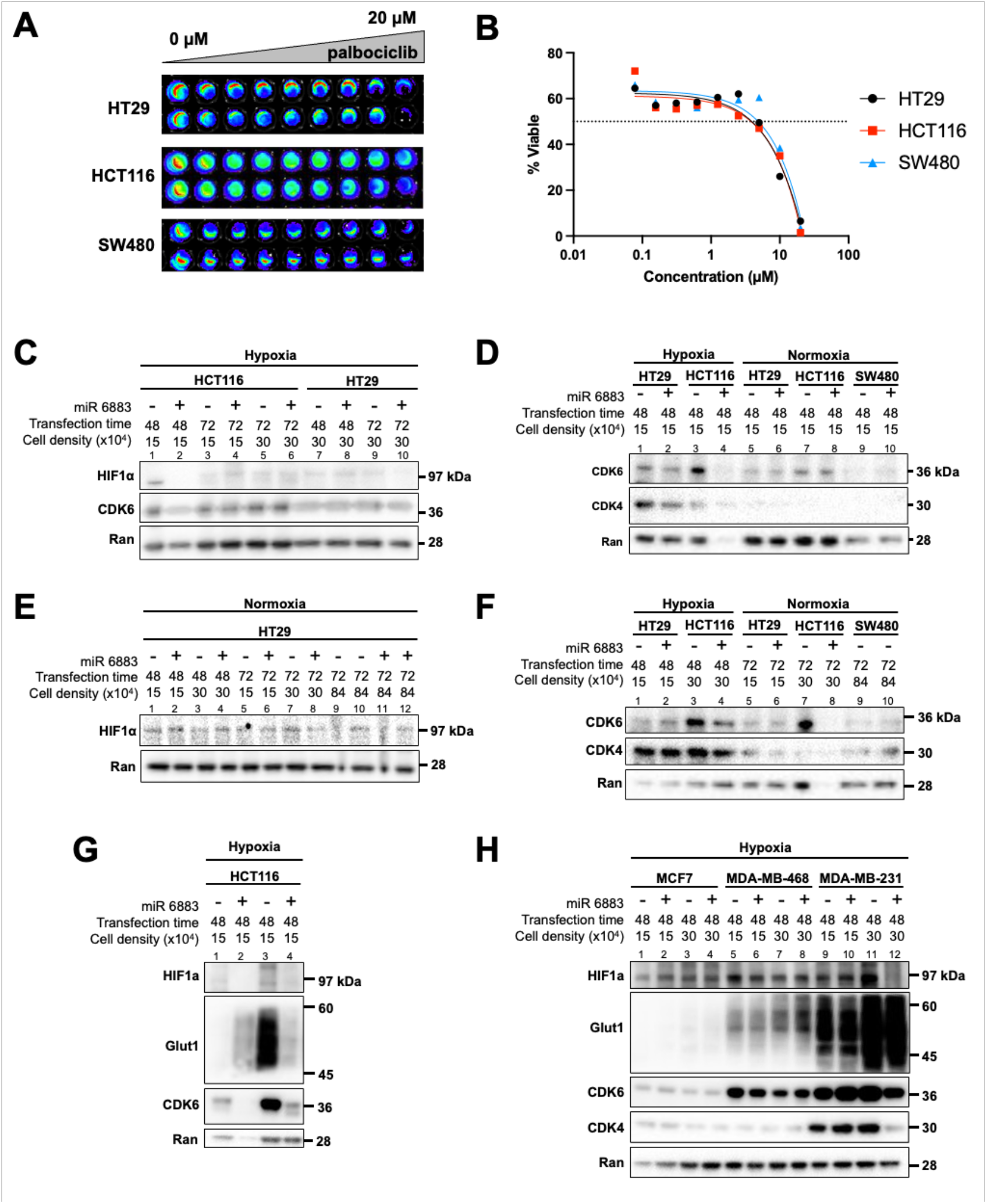

## Description

Colorectal Cancer (CRC) is characterized by the proliferation of abnormal cells in the colon and rectum. CRC originates from small polyps, which are often benign but can become large and cancerous. In 2023, there were an estimated 106,970 new cases of colon cancer and 46,050 new cases of rectal cancer in the United States, although recently incidence has been decreasing around 1% each year due to early screening, fecal immunochemical tests (FIT), and colonoscopies (*Key Statistics for Colorectal Cancer*, 2023; Roselló et al., 2019). Mortality is also decreasing and CRC-related deaths in 2023 are estimated to be 52,550 compared to 53,200 in 2020 (*Colorectal Cancer Facts & Figures 2020-2022*, 2020). However, CRC remains the fourth most diagnosed cancer and the second leading cause of cancer-related mortality globally. Colorectal cancer incidence has been rising over the last two decades among younger individuals for unclear reasons. Though treatments for CRC such as surgery, chemotherapy, targeted therapy, immunotherapy, and radiation can help to prolong patients’ lives, additional treatment options are needed for this deadly disease (*Colon cancer*). Excluding skin cancers, breast cancer is the most common cancer in women (*Key Statistics for Breast Cancer*) and has a 5-year survival rate of 30% once it spreads to distant organs (*Survival Rates for Breast Cancer*, 2023). Breast cancer is normally treated with surgery, chemotherapy, hormone therapy, or radiation but new treatments are needed to improve survival outcomes while avoiding treatment-related toxicities (*Breast cancer*).

Micro-RNAs (miR’s) are a group of non-coding RNAs that control coding genes (Mehrgou et al., 2021). miR’s function by binding to mRNA and preventing their translation. miR’s can bind oncogenes to suppress the initiation, progression, metastasis, and recurrence of CRC (To et al., 2018). Current clinical trials are investigating the efficacy of miR’s in various types of cancer such as lymphoma, mycosis fungoides, melanoma, lung cancer, liver cancer, and myeloma (Menon et al., 2022; O’Neill, 2016; Zhang et al., 2021).

The tumor microenvironment often becomes hypoxic as a result of growth of the tumor that outstrips its blood supply leading to inadequate oxygen delivery also because of atypical tumor blood vessels and overconsumption of oxygen within the tumor (Li et al., 2021; Muz et al., 2015). When cancer cells detect a hypoxic environment, hypoxia inducible factor 1 alpha (HIF1α) is stabilized (Cao et al., 2009). Hypoxic conditions and HIF1α signaling have a pro-angiogenic effect and contribute to the formation of endothelial cells, which improves the delivery of oxygen and nutrients to the tumor (Krock et al., 2011).

Cyclin-dependent kinases (CDKs) play a crucial role in cell cycle regulation. They bind cyclins, allowing transitions between G1, S, G2, and mitosis. Dysregulation of CDK activity contributes to the development of cancer. CDK4/6 inhibitors block cancer cell proliferation by inducing cell cycle arrest at G1 (Goel et al., 2018). CDK4/6 is targeted by several FDA-approved cancer therapeutics including palbociclib, ribociclib, and abemaciclib. Previously, our lab has shown that CDK4/6 inhibition destabilizes HIF1α in CRC cells (Zhao et al., 2021) and that miR-6883 can inhibit CDK4/6 (Lulla et al., 2017). We hypothesized that miR-6883 downregulates CDK4/6 and HIF1α in CRC and BC cells.

We confirmed that CRC cells are sensitive to CDK4/6 downregulation. HCT116, HT29, and SW480 cells were treated with increasing doses of the CDK4/6 inhibitor palbociclib for 72 hours and a CellTiter-Glo assay was used to measure cell viability. Bioluminescent imaging (panel A) and analysis in GraphPad Prism (panel B) revealed that CRC cells are sensitive to palbociclib (IC50 = ∼8 µM), providing a rationale to test the CDK4/6-targeting miR-6883 as a potential therapeutic agent for this disease. As CDK4/6 inhibitors are already used to treat patients with BC, it was not necessary to confirm their sensitivity to palbociclib.

We evaluated the effect of miR-6883 on CDK4/6 levels in CRC (HT29, HCT116) and BC (MCF7 and MDA-MB-231) cells. We transfected the tumor cells with miR-6883 using lipofectamine RNAiMAX and measured protein levels of CDK4/6 and HIF1α by western blot. After testing different cell densities and transfection times, we selected transfection conditions that successfully downregulated CDK4/6. In HCT116 cells, miR-6883 downregulation of CDK4/6 (panel C lanes 1-2, panel F lanes 3-4, panel G lanes 3-4) co-occurred with downregulation HIF1α (panel C lanes 1-2, panel G lanes 3-4) and HIF1α target gene Glut 1 (panel G lanes 3-4). Similar results were obtained with HT29 cells, in that miR-6883 downregulated CDK4/6 (panel D lanes 1-2) and HIF1α (panel E lanes 7-8). miR-6883 transfection was unsuccessful in SW480 CRC cells in the experiment shown (panel D lanes 9-10, panel F lanes 9-10). In MDA-MB-231 BC cells, transfection of miR-6883 in MDA-MB-231 caused downregulation of CDK4/6 which co-occurred with HIF1α downregulation (panel H lanes 11-12) with no effect of HIF1α target gene Glut1. Transfection of MCF7 (panel H, lanes 1-4) and MDA-MB-468 (panel H lanes 5-8) BC cells was unsuccessful in the experiments shown with no expected effect on HIF1α.

In summary, our results indicate the miR-6883 targets CDK4/6 in CRC (HCT116 and HT29) and BC (MDA-MB-231) cells. Downregulation of CDK4/6 co-occurred with downregulation of HIF1α, pointing to miR-6883 as a promising therapeutic for CRC and BC tumors with CDK4/6 and/or HIF1α overactivity. The proposed therapeutic strategy allows for miR-6883 mediated targeting of production CDK4/6 protein thereby reducing the pro-tumorigenic effects of HIF1α. The strategy also opens up a novel approach to address tumor progression and resistance to cancer therapy through either CDK4/6 or HIF1α.

## Methods

### Cell lines and culture conditions

CRC (HT29, HCT116, and SW480) and BC (MCF7, MDA-MB-468 and MDA-MB-231) cells were obtained from ATCC. HT29 and HCT116 cells were grown in McCoy’s 5A Medium supplemented with 10% FBS and 1% penicillin/streptomycin. MCF7 cells were grown in Minimum Essential Eagle Medium MEM 1X supplemented with 10% FBS and 1% penicillin/streptomycin. SW480, MDA-MB-468 and MDA-MB-231 cells were grown in DMEM media supplemented with 10% FBS, 1% sodium pyruvate, and 1% penicillin/streptomycin. All cells were incubated at 37°C, 5% CO_2_.

### CellTIterGlo Assay

CRC cells were plated at a density of 5,000 cells per well of a 96-well plate. Cells were treated with palbociclib using doses ranging from 0-20 μM. Cell viability was measured by adding CellTiter-Glo reagent followed by bioluminescence imaging.

### Western Blot

Cells were harvested and lysed using a RIPA buffer containing a protease inhibitor. Denaturing sample buffer was added, samples were boiled at 95°C for 10 minutes, and an equal amount of protein lysate was electrophoresed through 4-12% SDS-PAGE gels (Invitrogen) then transferred to PVDF membranes. The membrane was blocked with 5% milk in 1 × TTBS, incubated overnight in the appropriate primary antibody (HIF1α, pRb, Glut1, CDK4/6, or Ran), and incubated in the appropriate HRP-conjugated secondary antibody for two hours. The levels of antibody binding were detected using ECL western blotting detection reagent and the Syngene imaging system.

### Transfection

CRC cells (HT29, HCT116 and SW480) and BC cells (MCF-7, MDA-MB-468 and MDA-MB-231) were plated at various cell densities and incubated for at least 12 hours to allow cells to adhere to the plate. The cells were then transfected with 50 nmol/L miR-6883 mimic (Sigma-Aldrich (HMI2616) using Lipofectamine® RNAiMAX Reagent. The cells were incubated for either 48 or 72 hours under normoxia (∼20.9% O_2_) and hypoxia (<0.5% O_2_).

### miR-6883

miR-6883 mimic was purchased from Sigma-Aldrich (HMI2616). This double-stranded RNA molecule mimics endogenous mature miRNA when introduced into cells.

## Reagents

| **Antibody** | **Source** | **Manufacturer and catalog #** |
| --- | --- | --- |
| Anti-HIF1α | Mouse | Becton Dickinson 610958 |
| Anti-Rb | Mouse | Cell Signaling 9309 |
| Anti-pRb | Rabbit | Cell Signaling 8180 |
| Anti-Glut1 | Rabbit | Cell Signaling 73015 |
| Anti-CDK6 | Mouse | Cell Signaling 3136 |
| Anti-CDK4 | Rabbit | Cell Signaling 12790 |
| Anti-Ran | Mouse | Becton Dickinson 610341 |
| Anti-VEGF-A | Rabbit | Cell Signaling 50661 |
| Anti-mouse IgG | Goat | Thermo Fisher Scientific 31430 |
| Anti-rabbit IgG | Rabbit | Thermo Fisher Scientific 31460 |

## Acknowledgements

W.S.E-D. is supported by the Mencoff Family Professorship at Brown University. W.S.E-D. is an American Cancer Society Research Professor.

## Author Contributions

Conceptualization, W.S.E-D. Investigation, N.J-V., L.C., and W.S.E-D. Writing—original draft preparation, N.J-V. and L.C. Writing—review and editing, N.J-V., L.C., and W.S.E-D. Supervision, W.S.E-D. Project administration, W.S.E-D. Funding acquisition, W.S.E-D.

